# GLOBAL BIOGEOGRAPHY AND EVOLUTIONARY DRIVERS OF FERNS WITH CHLOROPHYLLOUS SPORES

**DOI:** 10.1101/2024.11.18.624067

**Authors:** Daniela Mellado-Mansilla, Patrick Weigelt, Michael Kessler, Dylan Craven, Gerhard Zotz, Holger Kreft

## Abstract

About 14% of all fern species have chlorophyllous spores. Unlike their counterparts, chlorophyllous spores lack dormancy and have a shorter viability (i.e. only a few days in some species). Such spores should limit long-distance dispersal and be more susceptible to harsher climatic conditions raising questions about the evolutionary and ecological advantages of this trait. This study aims to assess the global biogeography of chlorophyllous-spored ferns evaluating their underlying environmental and evolutionary drivers. We studied the global distribution of 10,995 fern taxa across 556 geographical regions, and assessed the association of the proportional representation of 1387 chlorophyllous-spored species (CSS) with environmental variables using generalized linear mixed models. To assess the influence of phylogenetic relationships on the distribution of this trait across the phylogeny and across geographic regions, we calculated the phylogenetic signal and phylogenetic distances among species in different assemblages. Species richness of chlorophyllous-spored ferns peaked in the tropics while their proportional representation was highest in temperate and island floras. The proportion of CSS was positively influenced by water availability, but negatively by variables associated with temperature. Spore type was strongly conserved phylogenetically, and CSS assemblages were phylogenetically clustered towards higher latitudes. Our study provides strong evidence that chlorophyllous spores do not limit the geographical distribution of fern species. Their latitudinal distribution patterns can be explain both for environmental and phylogenetic drives.

## Introduction

Ferns (i.e. class Polypodiopsida) are a diverse and one of the earliest groups of vascular plants with a evolutionary history of about 400 million years (Niklas et al. 1983, Testo and Sundue 2016). With approximately 10,578 species globally (PPG I 2016), ferns form the second largest group of vascular plants (Smith 1972), and play important ecological roles in many ecosystems (Hietz 1997, Ellwood and Foster 2004, Praptosuwiryo et al. 2019). However, fern diversity is increasingly threatened by global change drivers such as habitat loss and climate change. Although the consequences of global change on ferns are not well understood (Kessler and Kluge 2022), certain traits could play a critical role in determining the vulnerability of fern species to habitat changes.

Ferns exhibit distinct functional traits associated with each stage of their life cycle, namely unicellular airborne spores, gametophytes, and sporophytes. Each stage, along with its respective traits, plays a crucial role in the successful establishment of new fern populations. For instance, a single fern spore can disperse to new locations, where its germination results in a bisexual gametophyte capable of self-fertilization, which subsequently gives rise to the sporophyte stage (Haufler et al., 2016; Ranker et al., 1994; Smith, 1972). Therefore, the survival of spores during dispersal and their ability to germinate in new locations represent a critical bottleneck in the establishment of new fern populations. This study focuses on this initial step of the fern life cycle, examining the role of spore type in the global distribution of ferns.

Fern spores can be classified as chlorophyllous or non-chlorophyllous, depending on the presence or absence of chloroplasts when mature (Lloyd and Klekowski 1970, Sundue et al. 2011). Non-chlorophyllous spores can remain viable for extended periods, even years or decades, whereas chlorophyllous spores lack dormancy and typically retain viability for less than two months (Lloyd and Klekowski 1970). The differences in viability are mainly modulated by higher respiration rates and water loss due to the thinner walls of chlorophyllous spores, which could make them more vulnerable to harsh environmental conditions (Lloyd and Klekowski 1970). Despite clear differences in the viability of chlorophyllous and non-chlorophyllous spores, few studies have explored how this trait affects the biogeography of ferns (Dassler and Farrar 2001, Kessler 2002, Aldasoro et al. 2004) or which environmental variables influence the global distribution of chlorophyllous-spored species (hereafter CSS). Here, we fill this gap and explore the global distribution of CSS and determine its underlying environmental and evolutionary drivers.

It is estimated that approximately ∼1400 species (or 14%) of all extant species in the class Polypodiopsida have chlorophyllous spores (Mellado-Mansilla et al. 2022). Approximately 77% of CSS are epiphytes (1080 species), which represents ca. 37% of total epiphytic fern richness (Mellado-Mansilla et al. 2022). Additionally, c. 20% of the CSS are terrestrials and 3% are hemiepiphytes or climbers. Previous observations have linked the habits of CSS to more humid and stable climatic conditions. For instance, in temperate zones of the eastern United States, CSS are terrestrial and abundant in waterlogged soils and swamps (Sundue et al. 2011), whereas in tropical humid forests, where water availability is high, CSS are mostly epiphytes (Lloyd and Klekowski 1970). In line with this, the lack of dormancy in fern spores may have contributed to the success of CSS in environments characterized by consistent humidity where the likelihood of water loss is lower. Based on these observations, we hypothesized (H1) that the proportional representation of CSS in regional fern floras increases with high precipitation and low seasonality. Furthermore, we expect to find differences between the climatic variables driving the distribution of terrestrial and epiphytic CSS. Globally, the distribution of CSS should follow a similar latitudinal pattern as that of overall fern species richness, with higher species richness towards lower latitudes (Kreft et al. 2010, Weigand et al. 2020, Qian et al. 2023).

The relatively limited viability of chlorophyllous spores (Lloyd and Klekowski 1970) has prompted the idea that they may not be as effective in long-distance dispersal as non-chlorophyllous spores (Tryon 1970). However, this assumption was challenged by Dassler and Farrar (2001), who found that CSS belonging to Hymenophyllaceae and the grammitid group (Polypodiaceae) were over-represented on oceanic islands compared to the nearest mainland. Although germination of spores is the first bottleneck in the reproductive cycle of ferns, the presence of CSS on islands could be enhanced also by other traits such as the presence of gemmae in the gametophytes of some epiphytic CSS, which can facilitate vegetative reproduction and potentially enhance their dispersal capacity (Dassler and Farrar 2001). Furthermore, the autotrophic nature of chlorophyllous spores may confer an advantage in colonizing habitats with limited mycorrhizal associations (e.g. forest canopies, waterlogged soils, and oceanic islands) as germination is possible without fungal partners (Mellado-Mansilla et al. 2022). A similar advantage in the colonization of oceanic islands was also previously reported for angiosperm species independent on mycorrhizal associations (Delavaux et al. 2019). Overall, the autotrophic condition of chlorophyllous spores and the presence of gemmae in the gametophytes of CSS can collectively contribute to successful colonization and establishment on oceanic islands. Based on the traits associated with CSS and previous studies, we predict (H2) that the success in colonization and establishment of CSS in insular habitats will be higher than or similar to mainland regions. Therefore, we expect to find higher proportions of CSS on islands relative mainland regions.

Chlorophyllous spores have evolved several times independently, with four out of the six epiphytic lineages with major radiations occurring during the establishment of tropical rainforest in the Cenozoic having chlorophyllous spores (Schuettpelz and Pryer 2009, Sundue et al. 2015). CSS are ubiquitous in ancient families such as Hymenophyllaceae, Equisetaceae, Osmundaceae, Onocleaceae, and also in more recent lineages such as the grammitid group in Polypodiaceae. Furthermore, they are also found in some members of the genera *Aglaomorpha*, *Loxogramme*, *Platycerium*, *Pleurosoriopsis* (all Polypodiaceae), and *Elaphoglossum* (Dryopteridaceae), among others (Lloyd and Klekowski 1970, Sundue et al. 2011, Mellado-Mansilla et al. 2021). The arrangement of chlorophyllous spores in the fern phylogeny suggests a highly conserved trait, possibly indicative of phylogenetic niche conservatism. Recently, Qian et al. (2023) found that the global distribution of fern diversity can be mainly explained by the tropical niche conservatism hypothesis, according to which most fern species tend to conserve their ancestral ecological niche preferences (i.e. tropical niche) and only a few lineages evolved crucial traits to adapt to harsher extratropical environments (Wiens and Donoghue 2004). Consequently, phylogenetic relatedness among fern species assemblages increases toward higher latitudes (Wiens and Donoghue 2004, Qian et al. 2023). Here, we hypothesized (H3) that, coinciding with the general pattern observed for ferns, the phylogenetic relatedness of CSS assemblages increases towards higher latitudes, a pattern that supports the tropical niche conservatism hypothesis (Wiens and Donoghue 2004).

## Materials and methods

### Trait information

We used data on the presence and absence of chlorophyll in fern spores by Mellado-Mansilla et al. (2021). The total number of species with information on spore type was 2637 species (i.e. ca. 20 % of all fern species). We thus extrapolated this trait to all species of those genera in which only one spore type -either achlorophyllous or chlorophyllous - has been reported, whereas we did not do this in the genera *Aglaomorpha*, *Loxogramme*, *Platycerium*, *Pleopeltis*, *Pleurosoriopsis, Elaphoglossum* (Dryopteridaceae)*, Polytaenium (*Pteridaceae*), Lomaria* (Blechnaceae), and *Lomariopsis* (Lomariopsidaceae) because these contain both CSS and non-CSS spores. This increased the total number of fern species to be assessed almost four-fold, i.e. to 1387 ferns with chlorophyllous spores and 8278 ferns with achlorophyllous spores.

Since terrestrial and epiphytic ferns have different evolutionary histories (Schuettpelz and Pryer 2009; Sundue et al. 2015) and respond differently to abiotic variables that determine their distribution (Taylor et al. 2021), we tested our hypotheses for the overall CSS, and also for epiphytic and terrestrial CSS separately. Information on habit (terrestrial and epiphytic) was obtained from Zotz et al. (2021), field observations and bibliographic research from part of the authors of this study (a detailed list can be found in Mellado-Mansilla et al. 2022). Species nomenclature followed Hassler (2020). We excluded hybrids and the non-native range of species.

### Global distribution of fern species

The geographical distribution of each fern species was obtained from the Global Inventory of Floras and Traits database (GIFT version 2.1; (Weigelt et al. 2020)) based on the World Ferns database (Hassler 2020). World Ferns is the most comprehensive database on fern taxonomy and distributions assembled over the last 40 years and based on phylogenetic sequences and floristic checklists. For detailed information on the specific literature used to compile the fern flora of each geographic region see Hassler 2020. We used the information available for 556 non-overlapping geographical entities at the global scale that comprised countries or their political sub-units for countries with wide climatic variation, and also islands (178), and archipelagos (20). We excluded Russia from our analyses because of its large climatic and habitat heterogeneity without information on fern species richness at a smaller spatial scale.

### Environmental predictors

We used environmental variables previously reported as relevant to explain the global distribution of pteridophytes (Kreft et al. 2010, Weigand et al. 2020), including climatic variables, habitat heterogeneity, and botanical realms (Table 1). We included different precipitation-related variables due to the strong relationship between water availability and the germination of CSS described in previous studies (Ballesteros et al. 2011, López-Pozo et al. 2019a, b). We also included spatial entity (island/continent) as fixed effects due to the differences in richness of CSS in both geographical entities reported previously (Dassler and Farrar 2001, Aldasoro et al. 2004). For most variables, except elevational range, we used three summary statistics (mean, minimum, and maximum) per geographical entity. Potential evapotranspiration, mean annual cloud frequency, actual evapotranspiration, and habitat homogeneity were obtained from Zomer et al. (2008), Wilson and Jetz, (2016), Trabucco and Zomer (2010), and Tuanmu and Jetz (2015), respectively. All other environmental variables were obtained from CHELSA V.1.2 (Karger et al. 2017) and aggregated at a resolution of 30 arc-seconds for analyses.

**Table 1.** Single-predictor generalized linear mixed models (GLMMs) of the proportion of ferns with CS of ferns overall, the proportion of epiphytic ferns with CS, and the proportion of terrestrial ferns with CS per each geographic entity Each model included the random nested effect of realms and biomes. The r^2^s used to construct the multi-predictor final GLMMs was the marginal r^2^. All predictor variables were transformed. Sum. stat.: summary statistic.

|  | Overall |  | Epiphytes |  | Terrestrials |  |
| --- | --- | --- | --- | --- | --- | --- |
| Predictors | Sum. Stat. | $r^2$ | Sum. Stat. | $r^2$ | Sum. Stat. | $r^2$ |
| Potential evapotranspiration | max | 0.52 | max | 0.42 | max | 0.46 |
| Annual temperature | max | 0.47 | max | 0.33 | max | 0.61 |
| Prec. seasonality | mean | 0.30 | mean | 0.33 | max | 0.06 |
| Actual evapotranspiration | max | 0.15 | max | 0.18 | max | 0.46 |
| Mean annual cloud frequency | mean | 0.13 | max | 0.13 | min | 0.04 |
| Entity class (mainland) |  | 0.10 |  | 0.25 |  | .002 |
| Temperature annual range | max | 0.10 | min | 0.03 | min | 0.06 |
| Prec. driest month | max | 0.09 | max | 0.08 | min | 0.02 |
| Habitat homogeneity | mean | 0.07 | mean | 0.05 | mean | 0.01 |
| Elevational range |  | 0.06 | - | .002 | - | 0.01 |
| Prec. warmest quarter | max | 0.06 | max | 0.06 | max | 0.17 |
| Annual precipitation | mean | 0.03 | max | 0.01 | max | 0.04 |

### Environment-diversity relationships

We used Generalized Linear Mixed Models (GLMMs) to analyze the influence of the environmental variables on the proportional representation of CSS (H1). To account for spatial autocorrelation in our data we used realms as a random effect (1|REALM). We performed one GLMM for each response variable: a) the proportion of the overall CSS based on the total species richness of ferns, b) the epiphytic proportion of CSS based on the total richness of epiphytic species, and c) the terrestrial proportion of CSS based on the total species richness of terrestrial ferns. For all proportions, total fern species richness was the number of all fern species, including those with no data on spore type.

Since we used three summary statistics for each environmental variable, we started by testing the influence of each one of these variables on the proportions of CSS. We fitted single-predictor GLMMs and ranked variables according to the highest r^2^, and chose summary statistics with the highest r^2^ of the three statistic measurements. To avoid collinearity in our final models, we kept the variables with the highest r^2^ and discarded all variables correlated with them with a Pearson coefficient > 0.5. Second, we constructed GLMMs with the selected variables for each response variable. We used stepwise model selection to identify the best model based on the Akaike information criterion (AIC). Finally, we evaluated if spatial autocorrelation was present in the residuals of our final GLMMs performing a Moran’s I test with the function ‘testSpatialAutocorrelation’ of the Dharma R package (Hartig 2020). We centered and scaled all predictor variables (zero mean, one standard deviation). All GLMMs were performed with the R package glmmTMB (Brooks et al. 2017) using a beta distribution with a logit-link function.

### Phylogenetic metrics and analyses

To determine how the phylogenetic relationships may affect the geographic distribution of the CSS, we used the time-calibrated megaphylogeny of pteridophytes built by Hernández-Rojas et al. (2021) that includes around 50% of the extant fern species and was constructed using molecular data from seven chloroplast regions (atpΑ, atpΒ, matK, rbcL, rpl16, rps4, and trnL-trnF) and 26 fossils (Testo and Sundue 2016, Hernández-Rojas et al. 2021). The tree included 543 species with chlorophyllous spores and 2796 with non-chlorophyllous spores.

First, we determined how conserved is the spore type in the fern phylogeny. To this end, we calculated the phylogenetic signal of the spore type using the function ‘phylo.d’ of the R package Caper (Orme 2018). The D-statistic was designed to evaluate the phylogenetic signal in binary traits and compares the observed D values to simulated D values of the traits evolving under a Brownian model (D ≤ 0, strong phylogenetic signal) and under a model of random evolution (D = 1, no signal) (Fritz and Purvis 2010). The function returns p-values of testing whether D is statistically significantly different from 0 and whether is significantly different from 1. Non-significant p-values of D being different from 0 indicate that the trait is strongly phylogenetically conserved.

Second, to determine the phylogenetic relatedness of the CSS assemblages per geographic entity, we calculated the mean pairwise distance (MPD) and mean nearest taxon distance (MNTD) of each regional fern flora. MPD corresponds to the mean phylogenetic distance among all pairs of species within a region and emphasizes deep phylogenetic relationships, whereas MNTD corresponds to the mean distance between each one of the species within a region but considering its closest relative and emphasizes shallower phylogenetic relationships (Webb et al. 2002). Although MPD and MNTD are independent of species richness (De Bello et al. 2016), their variances can vary with sample size (Swenson et al. 2006). We thus used in all our analyses their standardized effect size by comparing the observed values to those expected under a null model. Our null models consisted of 999 randomizations of the phylogeny tip labels (i.e. species) in each geographical unit. Negative values of MPD and MNTD indicate that species in the assemblage are phylogenetically clustered (i.e. species within the assemblage are more closely related), while positive values indicate they are overdispersed (i.e. species are more distantly related). We calculated MPD and MNTD for all CSS and separately for epiphytic and terrestrial CSS. All calculations were performed using the functions ses.mpd and ses.mntd from the R package Picante (Kembel et al. 2010).

To test the tropical niche conservatism hypothesis (H3), we fitted linear mixed effects models with a similar structure as described above but using the standardized size effect of MPD and MNTD of CSS (overall, epiphytic, and terrestrials) as response variables. Since values of both response variables followed a normal distribution, we used the Gaussian family to perform GLMMs using the function lmer from the R package lme4 (Bates et al. 2009). All residuals and assumptions of our models were assessed using the R package performance (Lüdecke et al. 2021).

## Results

### Global distribution of chlorophyllous-spored species

CSS occurred in almost all geographic regions but with low species richness in arid regions and in the Arctic (<5 CSS species per geographical unit). Species richness of CSS peaked at lower latitudes, with marked hotspots in Central America and Southeast Asia (Fig. 1a, b; 267 species in Colombia, 223 species in New Guinea, and 212 species in Ecuador). Species richness in southern temperate zones was moderately higher (25-50 species) than in the northern temperate zones (5-10 species).

**Figure 1.**
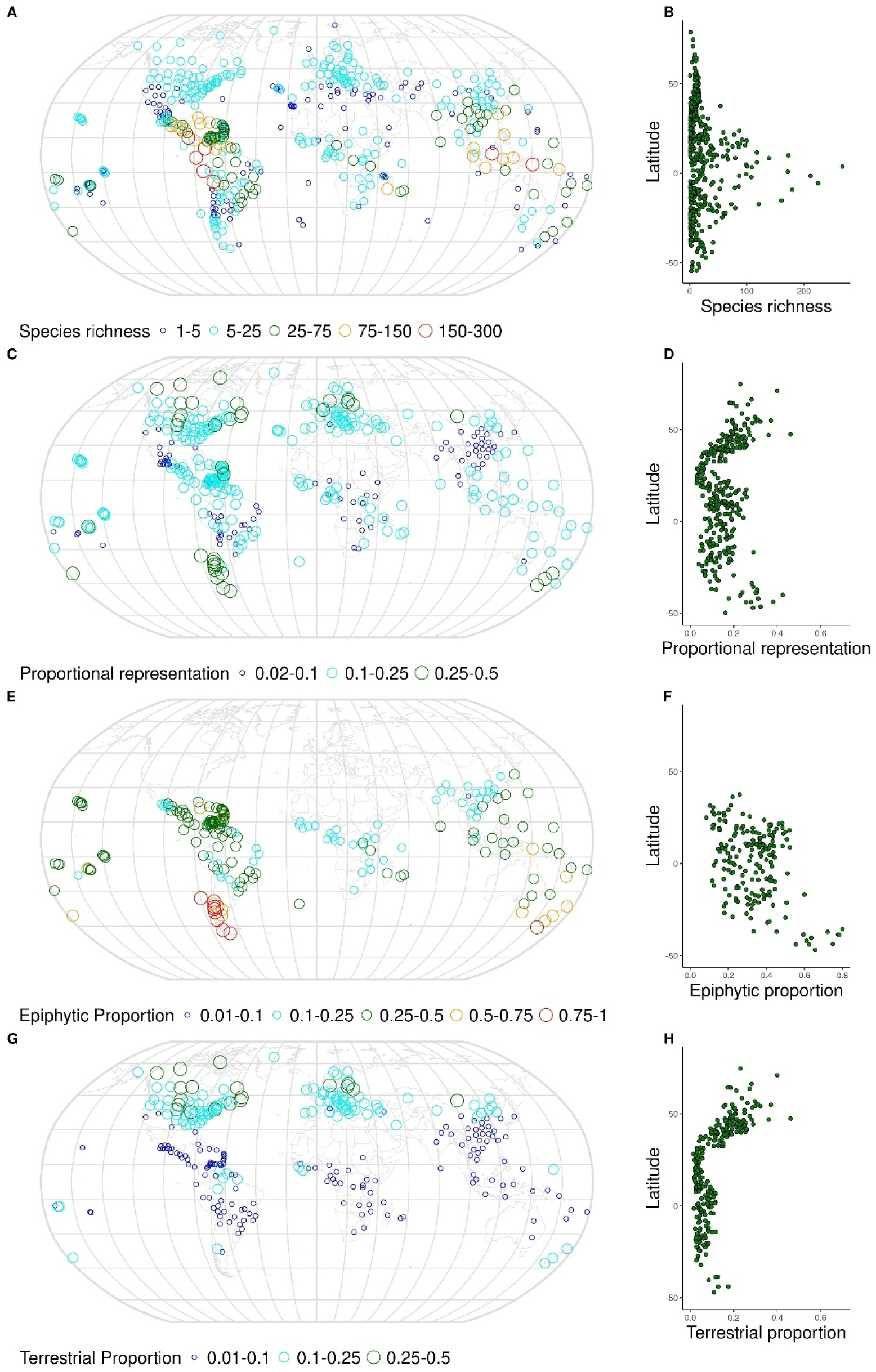
Global distribution of ferns with chlorophyllous spores. Only proportions of geographic entities with a richness >5 CSS are mapped.

Proportions of CSS were highest in the temperate zones of both hemispheres (25-50%) (Fig. 1c, d). The proportion of epiphytic CSS peaked in southern temperate zones, i.e. in southern Chile and Argentina, and New Zealand (60-86%) (Fig. 1e, f), including islands such as those belonging to the Juan Fernandez Archipelago (82%, 14 species) and Campbell Island (80%, 5 species). Conversely, the highest proportions of terrestrial CSS occurred in the northern temperate and subpolar zones, especially in northern USA and Svalbard (40-44%) (Fig. 1g, h).

When comparing the species richness and proportional representation of CSS between mainland regions and islands, we found no statistical difference in species richness (26.7 ± 29.4 (mean ± SD) vs. 24.6 ± 34.8, p<0.001, Kruskal-Wallis test), but the proportion of CSS was significantly higher on islands (0.18 ± 0.12 vs. 0.12 ± 0.07, p < 0.02, Kruskal-Wallis test) (Fig. 2b). Likewise, epiphytic species richness did not differ between mainland regions and islands (18.2 ± 28.4 vs. 17.7 ± 24.4, p = 0.7), but the proportional representation of epiphytic CSS was significantly higher on islands (0.40 ± 0.22 vs. 0.27 ± 0.19, p < 0.0001) (Fig. 2d). In contrast, the species richness of terrestrial CSS was higher on mainland regions (7.7 ± 5.8 vs. 5.96 ± 6.22, p < 0.0001) (Fig. 2e).

**Figure 2.**
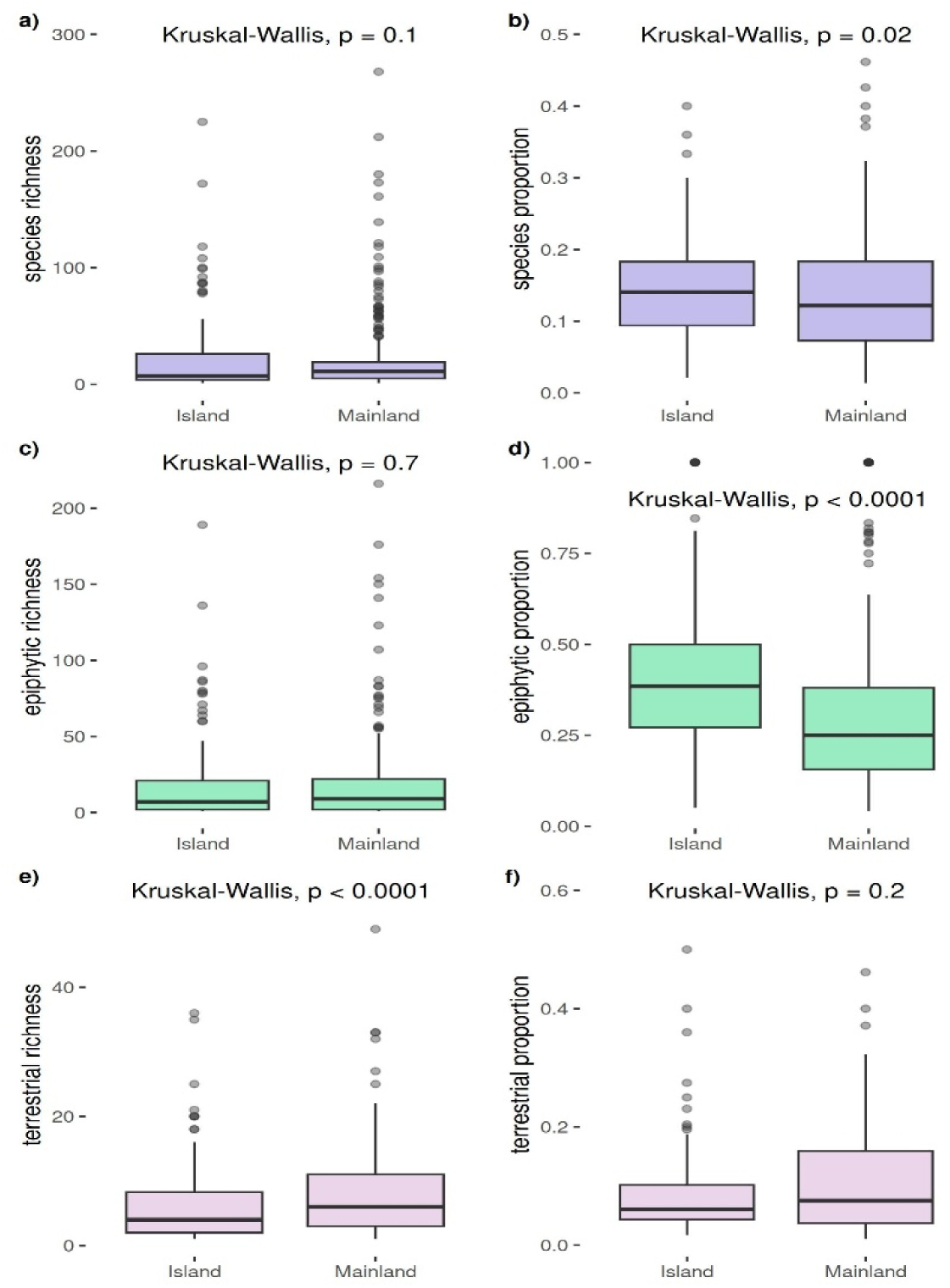
Species richness and proportional representation of CSS on islands and in mainlands.

### Abiotic drivers of CSS distribution

The most parsimonious model for the proportion of species richness of all CSS accounted for 66% of the variance including the random effect of realm (marginal r^2^ = 0.61). The proportional representation of all CSS increased with mean annual precipitation (0.12 ± 0.03, p < 0.001), but showed a negative association with maximum annual temperature (−0.39 ± 0.04, p < 0.001) and with maximum annual temperature range (−0.12 ± 0.06, p < 0.005) (Fig. 3). The final model of the proportion of epiphytic CSS had a total explanatory power of 94% (marginal r^2^ = 0.71) and the predictor variables with the strongest effects were mean annual temperature (−0.64 ± 0.08, p < 0.001) and mean cloud frequency (0.47± 0.06, p < 0.001). The final model for the proportion of terrestrial CSS explained a total of 87% of the model variance (marginal r^2^ = 0.85), and the most important variables were mean annual temperature (−0.44 ± 0.03, p < 0.001) and precipitation of the warmest quarter (−0.18 ± 0.04, p < 0.001) (Fig. 3). Finally, the residuals of the three final models exhibited a minor but not significant amount of spatial autocorrelation (Moran’s I: Overall = 0.03, epiphytes = 0.12, terrestrials = −0.22; all p > 0.05).

**Figure 3.**
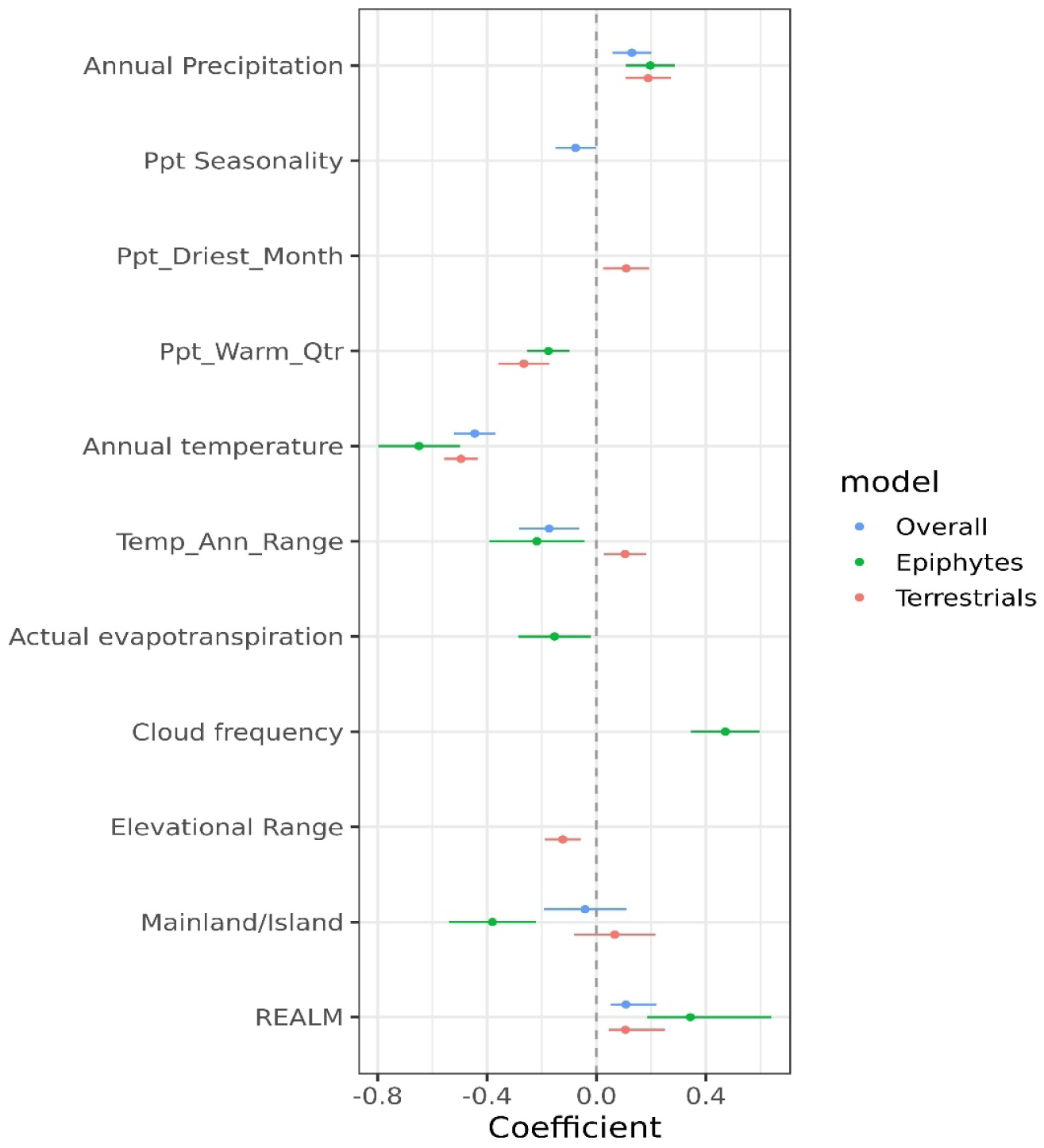
Coefficients plot of the three final GLMMs for chlorophyllous-spored species proportions. Overall: the overall proportional representation of ferns with chlorophyllous spores. Epiphytes and Terrestrials include the proportion of species richness with CS growing as epiphytes or terrestrials in each case.

### Evolutionary drivers of CSS distribution

As expected, spore type was strongly conserved phylogenetically (D = −0.36, p = 1). Regarding the phylogenetic indexes, MPD of terrestrial CSS was negative (i.e. clustered) in tropical and boreal zones (Fig. 4). MPD of epiphytic CSS were closer to 0 along the latitudinal gradient with negative values in Austral zones (Fig. 4c, d). MPD values of all CSS reflected both epiphytic and terrestrial patterns: most values were close to 0, with positive values in Asia and negative values in the Austral zones. Overall, MNTD values followed similar patterns to those of MPD and CSS habits (Fig. 5). Correlations between MPD and the richness proportion of CSS were weakly negative for the three groups (R= −0.26, −0.14, −0.31, p < 0.05, overall, epiphytic, and terrestrial respectively), indicating that higher proportions of CSS were weakly associated with phylogenetic clustering or dispersion (Fig. 6). Regarding MNTD, the correlation between epiphytic CSS proportion and MNTD was slightly stronger (−0.19, p < 0.005) than for the MPD but showed a positive relationship for the overall proportion (0.1, p < 0.05). However, MNTD was not correlated with terrestrial CSS (0.028, p > 0.05) (Fig. 6).

**Figure 4.**
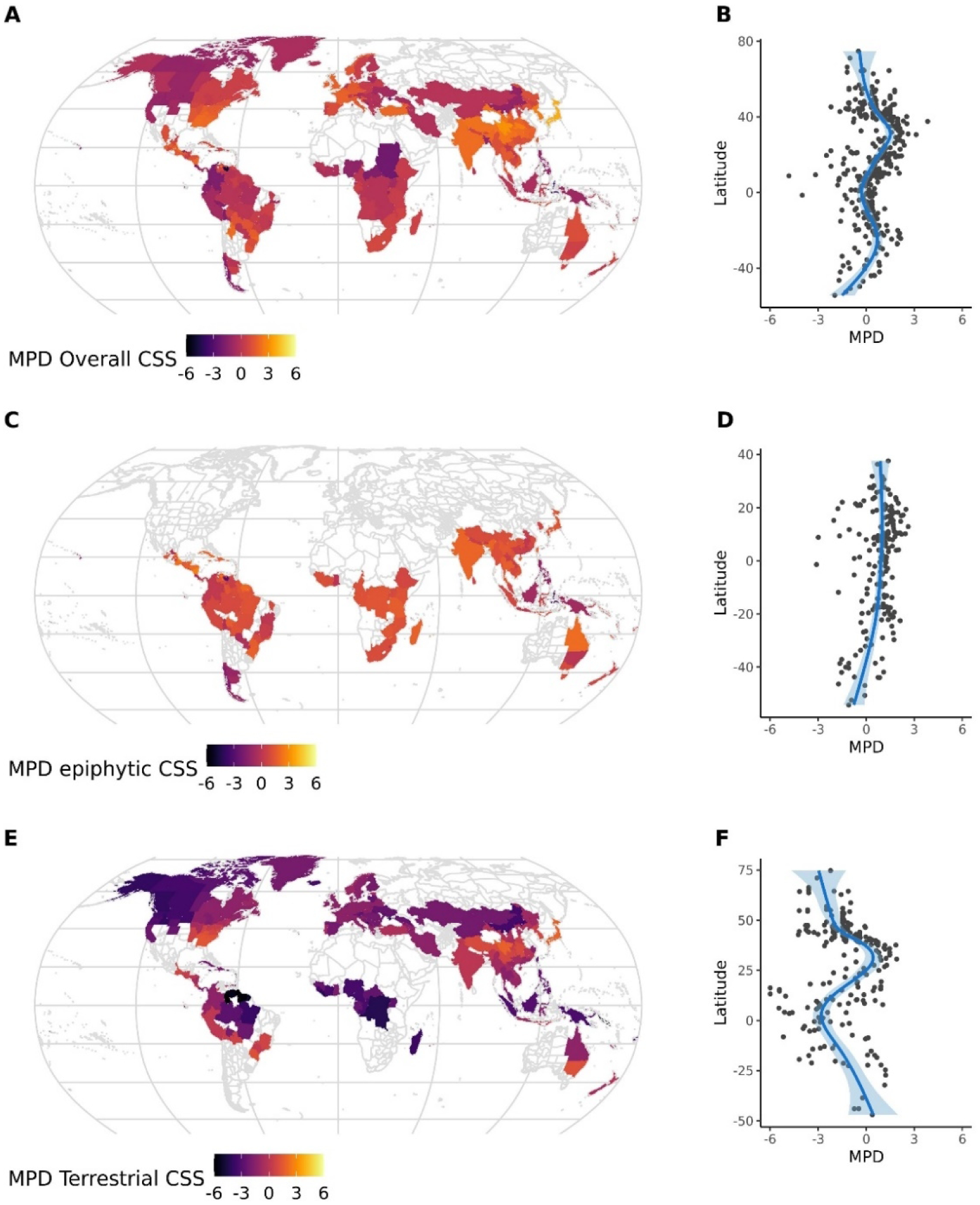
Mean pairwise distance (standardized effect size) globally for the overall proportion of CSS, the epiphytic proportion of CSS, and the terrestrial proportion of CSS. Negative values of MPD indicate phylogenetic clustering among species, while positive values indicate over-dispersion.

**Figure 5.**
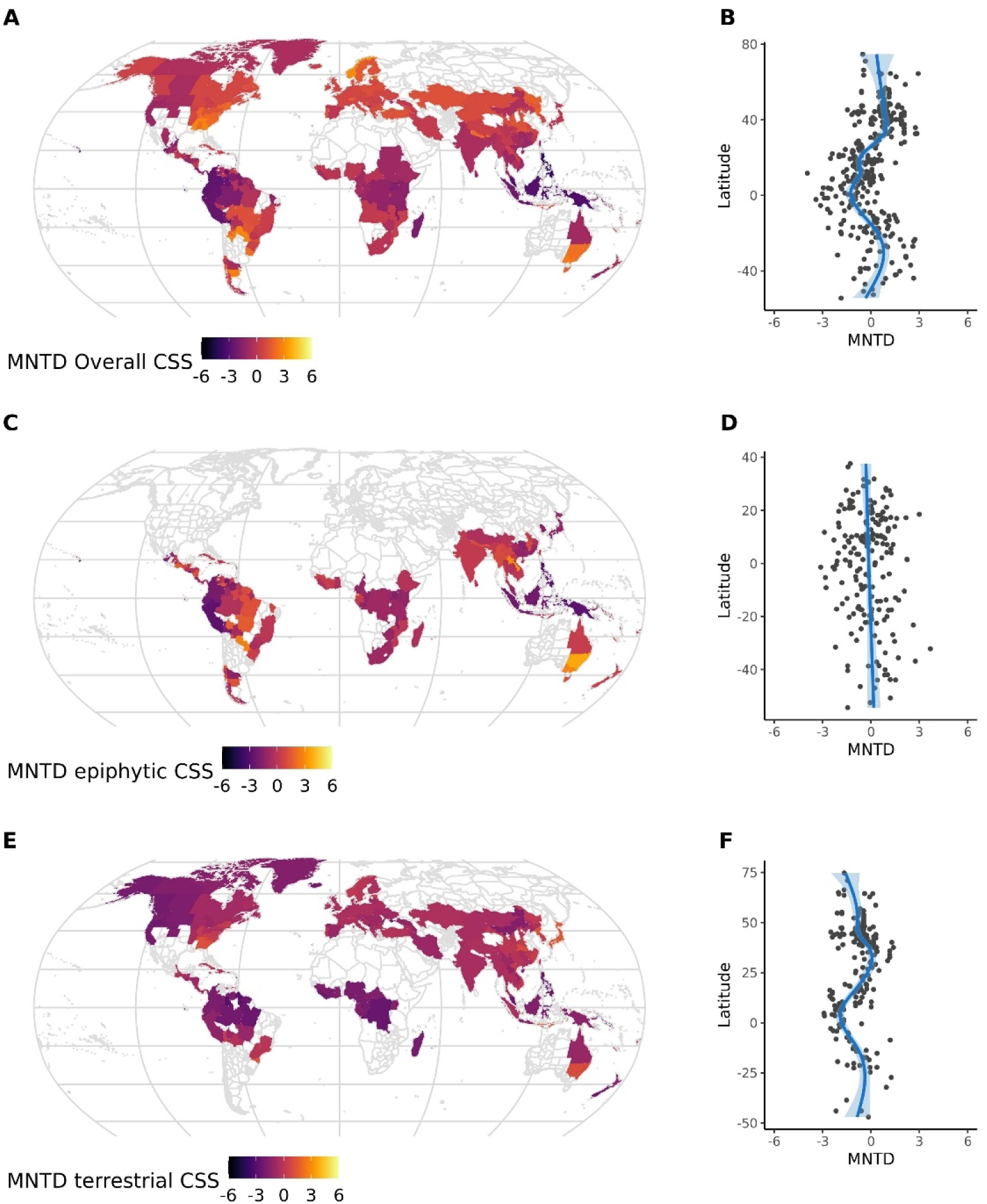
Mean nearest taxon distance (standardized effect size) for the overall proportion of CSS, the epiphytic proportion of CSS, and the terrestrial proportion of CSS. Negative values of MNTD indicate phylogenetic clustering among species, while positive values indicates overdispersion.

**Figure 6.**
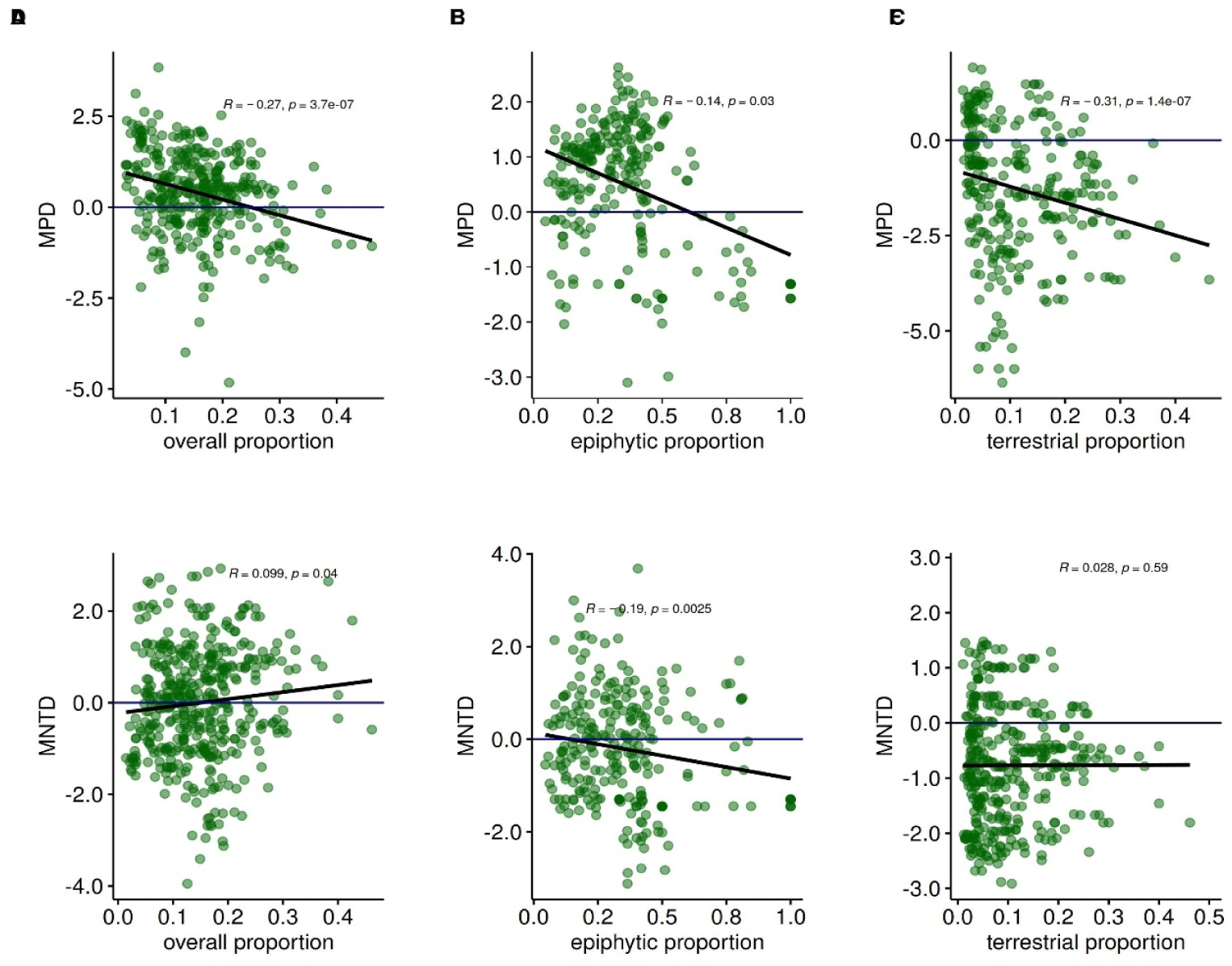
Correlations between the standardize effect sizes of mean pairwise distance (MPD), mean nearest taxon (MNTD), and the richness proportions of CSS. R= Spearman correlation coefficient, p= p-Value.

Environmental variables explained 34%, 27%, and 17% of the variance for the MPD of overall, epiphytic, and terrestrial CSS groups (marginal r² = 0.17, 0.22, and 0.15, respectively) (Fig. 7a). For both the overall and the epiphytic CSS, the environmental variables with the strongest effect on MPD were maximum annual temperature (overall: 0.30 ± 0.08, p < 0.005; epiphytes: 0.59 ± 0.14, p < 0.001) and mean precipitation of the warmest quarter (Overall: 0.31 ± 0.07, p < 0.005; epiphytes: 0.21 ± 0.07, p < 0.005). Regarding terrestrial CSS, maximum cloud frequency (−0.69 ± 0.16, p < 0.001) and maximum precipitation of the warmest quarter (0.33 ± 0.09, p < 0.001) were the variables with the strongest effect. Furthermore, environmental variables explained 31% of variation in MNTD of overall CSS (marginal r² = 0.31), and 27% of variation in MNTD of both epiphytic and terrestrial CSS (marginal r² = 0.23 and 0.19, respectively) (Fig. 7b). For the MNTD of overall CSS, the maximum temperature of the driest month (−0.43 ± 0.08, p < 0.001) and minimum precipitation of the warmest quarter (0.31 ± 0.08, p < 0.001) were significant variables. In the case of epiphytic CSS, MNTD variation was strongly influenced by maximum precipitation of the driest month (−0.18 ± 0.08, p < 0.01) and maximum cloud frequency (−0.34 ± 0.20, p < 0.05). Regarding terrestrial CSS, MNTD variation was mostly influenced by the mean precipitation of the warmest quarter (0.29 ± 0.07, p < 0.001), and the mean temperature annual range (0.28 ± 0.10, p < 0.01) (Fig. 7b). We did not detect a significant amount of spatial autocorrelation in the residuals of either six final models (Moran’s I: Overall MPD = −0.22, epiphytes MPD = −0.46, terrestrial MPD = −0.05; Overall MNTD = −0.11, epiphytes MNTD = −0.42, terrestrials MNTD = −0.05; all p > 0.05).

**Figure 7.**
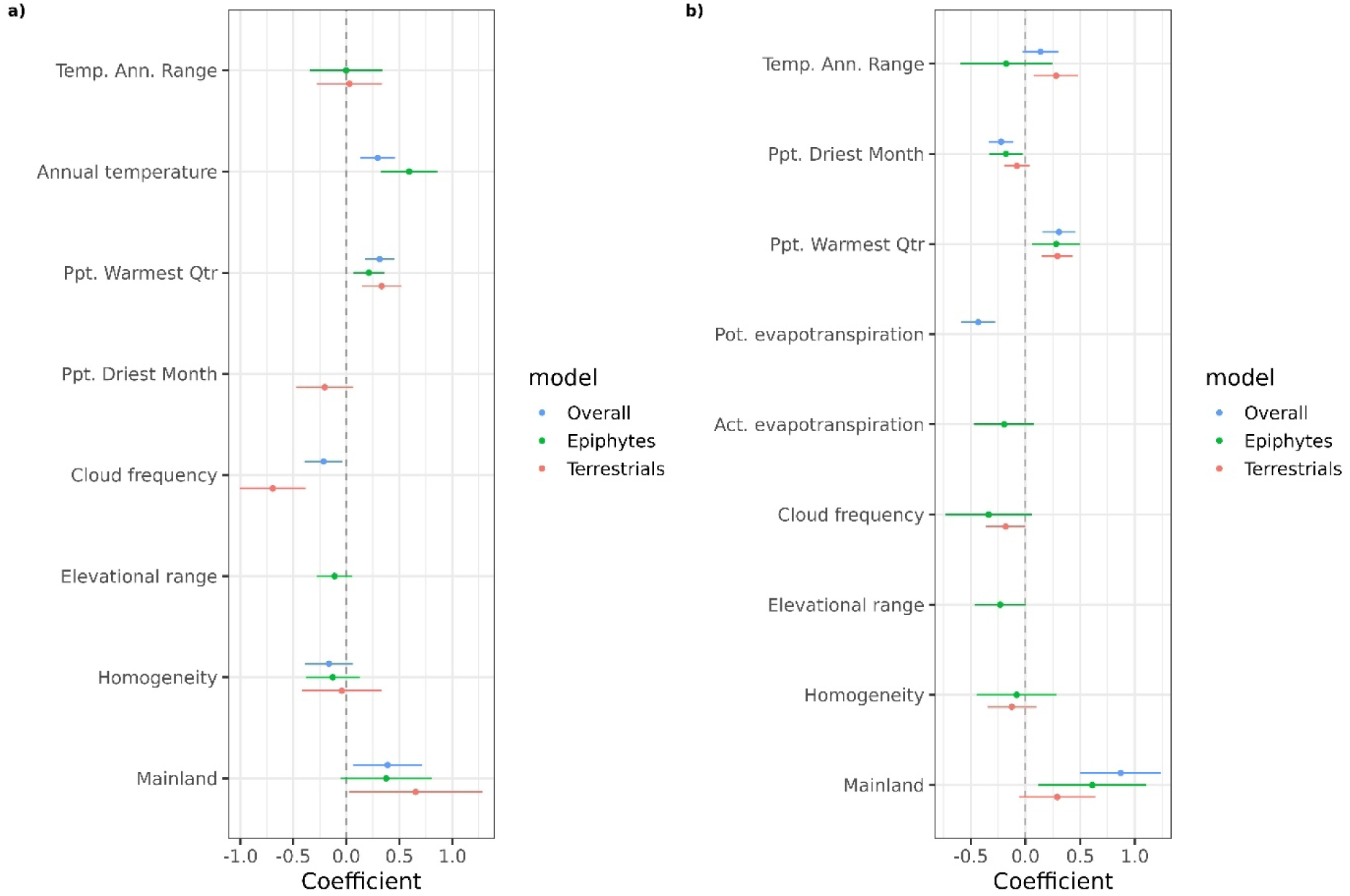
Coefficients plot of the final GLMMs for the standardize size effect of mean pairwise distance (a), and mean nearest taxon distance (b). Overall: the overall proportion of species richness of ferns with chlorophyllous spores. Epiphytes and Terrestrials include the proportion of species richness with CS growing as epiphytes or terrestrials in each case.

## Discussion

For the first time, we explored the environmental and evolutionary drivers of the spatial variation in the diversity and distribution of fern species with chlorophyllous spores. We found that the proportional representation of CSS is higher towards poles, and that, despite previous assumptions on their limited long-distance dispersal capacity, CSS are overrepresented on oceanic islands. Our results also showed that chlorophyllous spores not only have evolved multiple times in ferns, but it is also a highly conserved trait. Moreover, we found that the global distribution of CSS is the result of environmental and evolutionary drivers.

As in any macro-ecological study assessing the global distribution of a biological group, our study has some limitations. First, although our study is based on the most complete global dataset of fern distributions (Hassler 2020), aggregating environmental data across large spatial units could mask effects of microhabitat conditions in which CSS live. *Equisetum giganteum* L. living in the Atacama Desert is a good example to illustrate this point: when considering the climatic conditions of its distribution, we would be associating this species with arid conditions, even though it actually is associated with small wetlands. Considering these caveats, we can confirm that most of the patterns found in our study are strongly marked and therefore, using a different spatial scale, could make it possible to add other drivers of the distribution of CSS or, it could also emphasize the effect of the variables tested here.

Overall, we found a clear latitudinal gradient in the absolute number of CSS richness with higher peaks in equatorial zones as previously reported for global diversity of ferns (Kreft et al. 2010, Weigand et al. 2020, Suissa et al. 2021) (Fig. 1a, b), but strikingly, the proportional representation of CSS shows different patterns, peaking at higher latitudes where they tend to dominate fern floras (Fig. 1c, d). These patterns, however, are driven by different fern groups in both hemispheres. In the southern hemisphere, Hymenophyllaceae and grammitids dominate CSS in New Zealand and southern South America, while Equisetaceae, Osmundaceae, and Onocleaceae dominate in the north temperate zones. These differences are accompanied by a predominance of epiphytic CSS in the southern hemisphere, and of terrestrial ones in the northern hemisphere, which is in accordance with the higher overall vascular epiphytic richness in southern temperate forests compared to their northern counterparts as a result of several geological and climatic events such as the Last Maximum Glacial during the Pleistocene (Hinojosa and Villagran 1997, Villagran and Hinojosa 1997, Silander 2001, Zotz 2005, Taylor et al. 2021)

Our analyses of the proportional representation of CSS and environmental variables strongly support our hypothesis that their distribution is positively correlated with water-related factors and milder, less seasonal climates (Fig. 2). This result is further supported by the negative correlation between CSS and temperature-related variables, which is consistent with their prevalence in humid temperate regions. Furthermore, these patterns are mirrored by the different environmental variables influencing the CSS habits. For instance, terrestrial CSS are negatively associated with seasonality in precipitation, whereas epiphytic CSS are positively associated with the mean annual cloud frequency. While these patterns are consistent with the well-established idea that ferns generally depend on humid environmental conditions (Kessler et al. 2001, Kreft et al. 2010, Weigand et al. 2020), our study highlights that CSS are similarly dependent on water availability. Indeed, comparing our results to previous studies using similar environmental variables to explain the global distribution of ferns (Weigand et al. 2020), we found that the proportional representation CSS is mostly explained by precipitation-related variables. Yet, Qian et al. (2023) found that temperature-related variables were stronger drivers of phylogenetic diversity of polypods and old clades of ferns than precipitation-related variables. However, our results are probably influenced by the high proportion of epiphytic species with CS, most of which are highly dependent on constant water availability due to their poikilohydric condition, such as the species of the family Hymenophyllaceae (Proctor 2012).

One explanation for our findings could be that, unlike those of species with dormant spores, the spores of CSS germinate quickly and therefore establish more readily in areas with consistently high-water availability throughout the year. However, despite our finding that CSS diversity is highest in aseasonal, humid regions, there is growing evidence that spores of many CSS are surprisingly tolerant of low humidity during the germination stage (Ballesteros et al. 2017, López-Pozo et al. 2019a, b). Indeed, several observations have confirmed that CSS inhabiting the northern hemisphere (i.e. Osmundacea, Equisetaceae, Onocleaceae) released their spores during spring-summer seasons. Still, to our knowledge, there is no data on sporulation available for CSS from the southern hemisphere, and therefore the germination phenology and the characteristics of the growing season of these species are still unknown. This information could help to better understand what factors besides environmental ones could contribute to shaping where CSS occur.

Consistent with earlier reports at a regional scale (Dassler and Farrar 2001, Aldasoro et al. 2004, Sundue et al. 2014) and supporting our H2, our findings indicate that the over-representation of CSS on islands is a robust global phenomenon. Although chlorophyllous spores are short-lived, which may reduce survival during long-distance dispersal, previous studies indicate that the presence of the grammitid genus *Leucotrichum* and around 30 other grammitid species in Africa and Madagascar are the result of long-distance dispersal events from America (Rouhan 2012, Bauret et al. 2017), as are the species of grammitids inhabiting Hawaii (Sundue et al. 2014). Together, these findings show that the CSS can be successful in long-distance dispersal despite the short viability of their spores. However, the successful establishment of new CSS populations on islands depends not only on spore dispersal but also on other key traits. For instance, the advantage of CSS of being self-sufficient upon germination and less dependent on mycorrhizal fungi for gametophyte survival than non-CSS species (Mellado-Mansilla et al. 2022) could be key on oceanic islands where specialized mycorrhizal fungi may be underrepresented in the early stages of colonization because their plant partners are absent, leading to mutual inhibition for colonization, as proposed for angiosperms (Delavaux et al. 2019). Moreover, many CSS have gametophytes with gemmae, which allow for vegetative reproduction, and this could increase their success in long-distance colonization (Dassler and Farrar 1997). The advantages and disadvantages of chlorophyllous spores are most apparent on oceanic islands but may also apply to continental contexts as well.

Overall, our results provide evidence in support of our H3 on the phylogenetic conservatism of CSS. We found that spore type is a highly conserved trait in the phylogeny of ferns, i.e. species with chlorophyllous spores have retained this trait consistently over evolutionary time (Wiens et al. 2010). When exploring the correlations of MPD and MNTD with the proportional representation of CSS, we found a slight tendency for species to be more phylogenetically clustered where CSS are dominant. This suggests that CSS are more closely related in those assemblages than expected by chance and probably share similar traits. Finally, along the latitudinal gradient, we observed that CSS assemblages tend to be more clustered at higher latitudes and more overdispersed in tropical regions, which is especially pronounced among terrestrial species. Interestingly, certain terrestrial fern floras are overdispersed in southern Asia (Fig. 4), a pattern previously reported for old clades of ferns (Qian et al. 2023). On the other hand, epiphytic CSS assemblages mainly showed higher MPD and MNTD values across their distribution, but tended to cluster towards southern latitudes. These results suggest that the CSS distribution might be also driven by evolutionary processes where these species have developed and/or conserved traits - such as spore type - that have allowed them to persist out of the tropics.

We found shifts in the importance of drivers of phylogenetic structure over evolutionary time, as revealed by our analyses of spatial variation in MPD and MNTD of CSS. For instance, MPD revealed over-dispersion of terrestrial CSS in the northern hemisphere, suggesting that older lineages influence this relationship, presumably Equisetales, in which some of its members diversified under temperate conditions (Christenhusz et al. 2021). Moreover, our GLMMs showed that although both MNTD and MPD are influenced by similar environmental variables, MNTD variation was mostly explained by water-related variables, especially in epiphytic CSS (Fig. 7). This could be explained by the diversification of epiphytic clades such as grammitids and Hymenophyllaceae during the Cenozoic (∼100 mya) during the formation of tropical angiosperm forests (Schneider et al. 2004, Schuettpelz and Pryer 2009), as their current distribution is strongly driven by the presence of forests with high humidity. Furthermore, these differences can also be observed when considering the stronger effect of biogeographical realms on the variation of overall MPD (17%) compared to the MNTD (0%) in our GLMMs. This indicates that deeper evolutionary relationships among CSS have been strongly influenced by historical biogeographic factors, including regional differences in the effects of past major extinction events (Lehtonen et al. 2017), especially in Africa (Weigand et al. 2020), as well as the survival of ancestral lineages in refugia regions, especially in Malesia (Qian et al. 2023).

## Conclusions

Our findings reveal that fern species with chlorophyllous spores prevail in temperate floras where overall fern species richness decreases. This pattern is specially marked when comparing the habit that they use, with epiphytic CSS overrepresented in the southern temperate zones and terrestrial species overrepresented in the northern hemisphere. Another remarkable finding of this study is that CSS are overrepresented on islands globally, which refutes previous assumptions on their limited long-distance dispersal. Besides, we found that the presence of CSS is strongly associated with areas with high water availability and that their distribution is not only driven by environmental factors but can also be explained by evolutionary drivers resulting in the phylogenetic conservatism.

## Acknowledgments

This work was funded by the National Agency for Research and Development (ANID) / Scholarship Program / DOCTORADO BECAS CHILE/2018 – 72190330.

## Data availability

The R scripts used in this study are publicly accessible in the following repository: https://github.com/DMelladoMansilla

## References

Aldasoro, J. J., Cabezas, F. and Aedo, C. 2004. Diversity and distribution of ferns in sub-Saharan Africa, Madagascar and some islands of the South Atlantic. - Journal of Biogeography 31: 1579–1604.

Ballesteros, D., Estrelles, E., Walters, C. and Ibars, A. M. 2011. Effect of storage temperature on green spore longevity for the ferns *Equisetum ramosissimum* and *Osmunda regalis*. - Cryoletters 32: 89–98.

Ballesteros, D., Hill, L. M. and Walters, C. 2017. Variation of desiccation tolerance and longevity in fern spores. - Journal of Plant Physiology 211: 53–62.

Bates, D., Maechler, M. and Dai, B. 2009. lme4: Linear mixed-effects models using S4 classes. R package version 0.999375–31.

Bauret, L., Gaudeul, M., Sundue, M. A., Parris, B. S., Ranker, T. A., Rakotondrainibe, F., Hennequin, S., Ranaivo, J., Selosse, M.-A. and Rouhan, G. 2017. Madagascar sheds new light on the molecular systematics and biogeography of grammitid ferns: New unexpected lineages and numerous long-distance dispersal events. - Molecular Phylogenetics and Evolution 111: 1–17.

Brooks, M. E., Kristensen, K., van Benthem, K. J., Magnusson, A., Berg, C. W., Nielsen, A., Skaug, H. J., Machler, M. and Bolker, B. M. 2017. glmmTMB balances speed and flexibility among packages for zero-inflated generalized linear mixed modeling. - The R journal 9: 378–400.

Christenhusz, M. J. M., Chase, M. W., Fay, M. F., Hidalgo, O., Leitch, I. J., Pellicer, J. and Viruel, J. 2021. Biogeography and genome size evolution of the oldest extant vascular plant genus, *Equisetum* (Equisetaceae). - Annals of Botany 127: 681–695.

Dassler, C. L. and Farrar, D. R. 1997. Significance of form in fern gametophytes: clonal, gemmiferous gametophytes of *Callistopteris baueriana* (Hymenophyllaceae). - International Journal of Plant Sciences 158: 622–639.

Dassler, C. L. and Farrar, D. R. 2001. Significance of gametophyte form in long-distance colonization by tropical, epiphytic ferns. - Brittonia 53: 352–369.

De Bello, F., Carmona, C. P., Lepš, J., Szava-Kovats, R. and Pärtel, M. 2016. Functional diversity through the mean trait dissimilarity: resolving shortcomings with existing paradigms and algorithms. - Oecologia 180: 933–940.

Delavaux, C. S., Weigelt, P., Dawson, W., Duchicela, J., Essl, F., Van Kleunen, M., König, C., Pergl, J., Pyšek, P., Stein, A., Winter, M., Schultz, P., Kreft, H. and Bever, J. D. 2019. Mycorrhizal fungi influence global plant biogeography. - Nat Ecol Evol 3: 424–429.

Ellwood, M. D. F. and Foster, W. A. 2004. Doubling the estimate of invertebrate biomass in a rainforest canopy. - Nature 429: 549–551.

Fritz, S. A. and Purvis, A. 2010. Selectivity in mammalian extinction risk and threat types: a new measure of phylogenetic signal Strength in binary traits. - Conservation Biology 24: 1042–1051.

Hartig, F. 2020. DHARMa: residual diagnostics for hierarchical (multi-Level / mixed) regression models. - R package version 0.3. 1. in press.

Haufler, C.H., Pryer, K.M., Schuettpelz, E., Sessa, E.B., Farrar, D.R., Moran, R., Schneller, J.J., Watkins Jr, J.E. and Windham, M.D. 2016. Sex and the single gametophyte: Revising the homosporous vascular plant life cycle in light of contemporary research. BioScience 66: 928–937.

Hassler, M. 2020. World Ferns. Synonymic checklist and distribution of ferns and lycophytes of the world. Version 14.8.

Hernández-Rojas, A. C., Kluge, J., Noben, S., Reyes Chávez, J. D., Krömer, T., Carvajal-Hernández, C. I., Salazar, L. and Kessler, M. 2021. Phylogenetic diversity of ferns reveals different patterns of niche conservatism and habitat filtering between epiphytic and terrestrial assemblages. - Frontiers of Biogeography 13: e50023.

Hietz, P. 1997. Population dynamics of epiphytes in a Mexican humid montane forest. - The Journal of Ecology 85: 767.

Karger, D. N., Conrad, O., Böhner, J., Kawohl, T., Kreft, H., Soria-Auza, R. W., Zimmermann, N. E., Linder, H. P. and Kessler, M. 2017. Climatologies at high resolution for the earth’s land surface areas. - Sci Data 4: 170122.

Kembel, S. W., Cowan, P. D., Helmus, M. R., Cornwell, W. K., Morlon, H., Ackerly, D. D., Blomberg, S. P. and Webb, C. O. 2010. Picante: R tools for integrating phylogenies and ecology. - Bioinformatics 26: 1463–1464.

Kessler, M. 2002. Range size and its ecological correlates among the pteridophytes of Carrasco National Park, Bolivia. - Global Ecology and Biogeography 11: 89–102.

Kessler, M., Parris, B. S. and Kessler, E. 2001. A comparison of the tropical montane pteridophyte floras of Mount Kinabalu, Borneo, and Parque Nacional Carrasco, Bolivia. - Journal of Biogeography 28: 611–622.

Kessler, M. and Kluge, J. 2022. Mountain ferns: what determines their elevational ranges and how will they respond to climate change? - American Fern Journal 112: 285–302.

Kreft, H., Jetz, W., Mutke, J. and Barthlott, W. 2010. Contrasting environmental and regional effects on global pteridophyte and seed plant diversity. - Ecography 33: 408–419.

Lehtonen, S., Silvestro, D., Karger, D. N., Scotese, C., Tuomisto, H., Kessler, M., Peña, C., Wahlberg, N. and Antonelli, A. 2017. Environmentally driven extinction and opportunistic origination explain fern diversification patterns. - Sci Rep 7: 4831.

Lloyd, R. M. and Klekowski, E. J. 1970. Spore germination and viability in Pteridophyta: evolutionary significance of chlorophyllous spores. - Biotropica 2: 129–137.

López-Pozo, M., Gasulla, F., García-Plazaola, J. I. and Fernández-Marín, B. 2019a. Unraveling metabolic mechanisms behind chloroplast desiccation tolerance: chlorophyllous fern spore as a new promising unicellular model. - Plant Science 281: 251–260.

López-Pozo, M., Ballesteros, D., Laza, J. M., García-Plazaola, J. I. and Fernández-Marín, B. 2019b. Desiccation tolerance in chlorophyllous fern spores: are ecophysiological features related to environmental conditions? - Front. Plant Sci. 10: 1–15.

Lüdecke, D., Ben-Shachar, M., Patil, I., Waggoner, P. and Makowski, D. 2021. performance: An R package for assessment, comparison and testing of statistical models. - JOSS 6: 3139.

Mellado-Mansilla, D., Zotz, G., Kreft, H., Sundue, M. and Kessler, M. 2021. The taxonomic distribution of chlorophyllous spores in ferns: an update. - American Fern Journal 111: 150–156.

Mellado-Mansilla, D., Testo, W., Sundue, M. A., Zotz, G., Kreft, H., Coiro, M. and Kessler, M. 2022. The relationship between chlorophyllous spores and mycorrhizal associations in ferns: evidence from an evolutionary approach. - American J of Botany 109: 2068–2081.

Niklas, K. J., Tiffney, B. H. and Knoll, A. H. 1983. Patterns in vascular land plant diversification. - Nature 303: 614–616.

Orme, D. 2018. The caper package: comparative analysis of phylogenetics and evolution in R.: 1–36.

PPG I 2016. A community-derived classification for extant lycophytes and ferns. - J of Sytematics Evolution 54: 563–603.

Praptosuwiryo, T. N., Sumanto, S. and Cahyaningsih, R. 2019. Diversity and host preferences of ferns and lycopods epiphytes on palm trees. - Biodiversitas Journal of Biological Diversity 20: 3731–3740.

Proctor, M. C. F. 2012. Light and desiccation responses of some Hymenophyllaceae (filmy ferns) from Trinidad, Venezuela and New Zealand: poikilohydry in a light-limited but low evaporation ecological niche. - Annals of Botany 109: 1019–1026.

Qian, H., Kessler, M., Zhang, J., Jin, Y. and Jiang, M. 2023. Global patterns and climatic determinants of phylogenetic structure of regional fern floras. - New Phytologist 239: 415–428.

Ranker TA Floyd SK Trapp PG 1994 Multiple colonizations of *Asplenium adiantum nigrum* onto the Hawaiian archipelago. Evolution 48: 1364–1370.

Rouhan, G. 2012. Not so Neotropical after all: the grammitid fern genus *Leucotrichum* (Polypodiaceae) is also Paleotropical, as revealed by a new species from Madagascar. Systematic Botany 37: 331–338.

Schneider, H., Schuettpelz, E., Pryer, K. M., Cranfill, R., Magallón, S. and Lupia, R. 2004a. Ferns diversified in the shadow of angiosperms. - Nature 428: 553–557.

Schuettpelz, E. and Pryer, K. M. 2009. Evidence for a Cenozoic radiation of ferns in an angiosperm-dominated canopy. - Proc. Natl. Acad. Sci. U.S.A. 106: 11200–11205.

Silander, J. A. 2001. Temperate Forests. - In: Levin, S. A. (ed), Encyclopedia of Biodiversity (Second Edition). Academic Press, pp. 112–127.

Smith, A. R. 1972. Comparison of fern and flowering plant distributions with some evolutionary Interpretations for ferns. - Biotropica 4: 4–9.

Suissa, J. S., Sundue, M. A. and Testo, W. L. 2021. Mountains, climate and niche heterogeneity explain global patterns of fern diversity. - Journal of Biogeography 48: 1296–1308.

Sundue, M., Vasco, A. and Moran, R. C. 2011. Cryptochlorophyllous spores in ferns: nongreen spores that contain chlorophyll. - International Journal of Plant Sciences 172: 1110–1119.

Sundue, M., Parris, B. S., Ranker, T. A., Smith, A. R., Fujimoto, E. L., Zamora-Crosby, D., Morden, C. W., Chiou, W.-L., Chen, C.-W., Rouhan, G., Hirai, R. Y. and Prado, J. 2014. Global phylogeny and biogeography of grammitid ferns (Polypodiaceae). - Molecular Phylogenetics and Evolution 81: 195–206.

Sundue, M., Testo, W. L. and Ranker, T. A. 2015. Morphological innovation, ecological opportunity, and the radiation of a major vascular epiphyte lineage. - Evolution 69: 2482–2495.

Taylor, A., Keppel, G., Weigelt, P., Zotz, G. and Kreft, H. 2021. Functional traits are key to understanding orchid diversity on islands. - Ecography 44: 703–714.

Taylor, A., Zotz, G., Weigelt, P., Cai, L., Karger, D. N., König, C. and Kreft, H. 2022. Vascular epiphytes contribute disproportionately to global centres of plant diversity. - Global Ecology and Biogeography 31: 62–74.

Testo, W. and Sundue, M. 2016. A 4000-species dataset provides new insight into the evolution of ferns. - Molecular Phylogenetics and Evolution 105: 200–211.

Trabucco, A. and Zomer, R. J. 2010. Global soil water balance geospatial database. CGIAR Consortium for Spatial Information. - cgiarcsi. community: Published online, available from the CGIARCSI GeoPortal.

Tryon, R. 1970. Development and evolution of fern floras of oceanic islands. - Biotropica 2: 76–84.

Tuanmu, M.-N. and Jetz, W. 2015. A global, remote sensing-based characterization of terrestrial habitat heterogeneity for biodiversity and ecosystem modelling. - Global Ecology and Biogeography 24: 1329–1339.

Villagran, C. and Hinojosa, L. F. 1997. Historia de los bosques del sur de Sudamérica, II: Análisis fitogeográfico. - Revista chilena de historia natural 70: 241–267.

Webb, C. O., Ackerly, D. D., McPeek, M. A. and Donoghue, M. J. 2002. Phylogenies and community ecology. - Annu. Rev. Ecol. Syst. 33: 475–505.

Weigand, A., Abrahamczyk, S., Aubin, I., Bita-Nicolae, C., Bruelheide, H., I. Carvajal-Hernández, C., Cicuzza, D., Nascimento Da Costa, L. E., Csiky, J., Dengler, J., Gasper, A. L. D., Guerin, G. R., Haider, S., Hernández-Rojas, A., Jandt, U., Reyes-Chávez, J., Karger, D. N., Khine, P. K., Kluge, J., Krömer, T., Lehnert, M., Lenoir, J., Moulatlet, G. M., Aros-Mualin, D., Noben, S., Olivares, I., G. Quintanilla, L., Reich, P. B., Salazar, L., Silva-Mijangos, L., Tuomisto, H., Weigelt, P., Zuquim, G., Kreft, H. and Kessler, M. 2020. Global fern and lycophyte richness explained: How regional and local factors shape plot richness. - Journal of Biogeography 47: 59–71.

Weigelt, P., König, C. and Kreft, H. 2020. GIFT – A global inventory of floras and traits for macroecology and biogeography. - Journal of Biogeography 47: 16–43.

Wiens, J. J. and Donoghue, M. J. 2004. Historical biogeography, ecology and species richness. - Trends in Ecology & Evolution 19: 639–644.

Wiens, J. J., Ackerly, D. D., Allen, A. P., Anacker, B. L., Buckley, L. B., Cornell, H. V., Damschen, E. I., Jonathan Davies, T., Grytnes, J., Harrison, S. P., Hawkins, B. A., Holt, R. D., McCain, C. M. and Stephens, P. R. 2010. Niche conservatism as an emerging principle in ecology and conservation biology. - Ecology Letters 13: 1310–1324.

Wilson, A. M. and Jetz, W. 2016. Remotely sensed high-resolution global cloud dynamics for predicting ecosystem and biodiversity distributions (M Loreau, Ed.). - PLoS Biol 14: e1002415.

Zomer, R. J., Trabucco, A., Bossio, D. A. and Verchot, L. V. 2008. Climate change mitigation: A spatial analysis of global land suitability for clean development mechanism afforestation and reforestation. - Agriculture, Ecosystems & Environment 126: 67–80.

Zotz, G. 2005. Vascular epiphytes in the temperate zones–a review. - Plant Ecol 176: 173–183.

Zotz, G., Weigelt, P., Kessler, M., Kreft, H. and Taylor, A. 2021a. EpiList 1.0: a global checklist of vascular epiphytes. - Ecology 102: e03326.

